# Germline inherited small RNAs clear untranslated maternal mRNAs in *C. elegans* embryos

**DOI:** 10.1101/2020.02.03.919597

**Authors:** Piergiuseppe Quarato, Meetali Singh, Eric Cornes, Blaise Li, Loan Bourdon, Florian Mueller, Celine Didier, Germano Cecere

## Abstract

Inheritance and clearance of maternal mRNAs are two of the most critical events required for animal early embryonic development. However, the mechanisms regulating this process are still largely unknown. Here, we show that together with maternal mRNAs, *C. elegans* embryos inherit a complementary pool of small non-coding RNAs capable of triggering the cleavage and removal of hundreds of maternal mRNAs. These antisense small RNAs are loaded into the maternal catalytically-active Argonaute CSR-1 and cleave complementary mRNAs no longer engaged in translation in somatic blastomeres. Induced depletion of CSR-1 specifically during embryonic development leads to embryonic lethality in a slicer-dependent manner and impairs the degradation of CSR-1 embryonic mRNA targets. Given the conservation of Argonaute catalytic activity, we propose that a similar mechanism operates to clear maternal mRNAs during the maternal-to-zygotic transition across species.

## MAIN TEXT

The elimination of germline-produced mRNAs and proteins in somatic blastomeres and the concomitant activation of the zygotic genome—the maternal to zygotic transition (MZT)— is critical for embryogenesis^1^. Despite several mechanisms have been discovered to regulate the clearance of maternal mRNAs during the MZT, they only account for the regulation of a small fraction of decayed transcripts, suggesting that factors and pathways responsible for the decay of the majority of maternal mRNAs still remain elusive. In addition, distinct species can adopt different strategies to achieve the clearance of maternal mRNAs. This is why understanding how different species clear maternal mRNAs is fundamental to shed light on novel mechanisms regulating of one the most important and conserved transition happening at the beginning of embryogenesis. For instance, small RNAs, such as maternally-inherited PIWI-interacting RNAs (piRNAs) in *Drosophila*^2,3^ and zygotically-transcribed micro RNAs (miRNAs) in *Drosophila*, zebrafish and *Xenopus*^4–6^ have been shown to directly regulate the degradation of maternal mRNAs in embryos. However, miRNA functions are globally suppressed in mouse oocytes and zygotes^7,8^ and they do not appear to contribute to maternal mRNA clearance in *C. elegans* embryos^9^. This raises the question of whether other type of small RNAs might play a role in this process. In this regard, catalytically active Argonaute and endogenous small interfering RNAs (endo-siRNAs) have been proposed to play an essential role in oocytes and early embryos in place of miRNAs^10,11^. Therefore, in addition to the silencing of repetitive elements, the RNA interference (RNAi) pathway might regulate the removal of maternal mRNAs in animal embryos.

In the nematode *C. elegans*, two waves of maternal mRNA clearance have been documented so far. The first wave of maternal mRNA clearance, regulated by a consensus sequence in the 3’UTR, occurs during the transition from the oocyte to the 1-cell embryo^9^. A second wave of clearance occurs in the somatic blastomeres of the developing embryos^12,13^. However, to date no mechanisms are known to regulate this process in the embryo. Endo-siRNAs produced in the germline are known to be heritable and they can potentially regulate mRNA transcripts in the developing embryos. Two main endogenous small RNA pathways regulate heritable epigenetic processes in the *C. elegans* germline: (1) thousands of PIWI-interacting RNAs (piRNAs) that target foreign “non-self” mRNAs^14^, and (2) endogenous antisense small RNAs that target active “self” mRNAs^15,16^. These antisense small RNAs, called 22G-RNAs, are generated by the RNA-dependent RNA polymerase (RdRP) EGO-1 using target germline mRNAs as a template and are then loaded into the Argonaute CSR-1 (Chromosome-Segregation and RNAi deficient)^16,17^. CSR-1 22G-RNAs are thought to protect germline mRNAs from piRNA silencing and they are essential for fertility and embryonic development^15,18,19^. However, CSR-1 is also responsible for the majority of slicing activity in *C. elegans* extracts and is capable of cleaving complementary mRNAs *in vitro*^20^. Moreover, CSR-1 slicer activity is capable of fine-tuning some of its mRNA targets in the adult germline^21^. Therefore, whether CSR-1 can protect or degrade its germline mRNA targets is still an open question.

Here, we provide novel insight into the widespread process of maternal mRNA clearance in early developing embryos. We demonstrate that 22G-RNAs loaded into CSR-1 that are inherited from the maternal germline, trigger the cleavage and removal of complementary maternal mRNAs in somatic blastomeres during early embryogenesis. Whereas in the germline CSR-1 and its associated 22G-RNAs result in a modest fine-tuning of target mRNAs (Gerson-Gurwitz et al., 2016), in the embryo CSR-1 targets are cleared. We found that target mRNAs that decay due to CSR-1 are depleted of ribosomes in embryonic somatic cells, and we provide evidence that the translational status of these maternal mRNAs influences their decay. We employed the auxin-inducible degradation (AID) system^22^ to deplete CSR-1 specifically during the MZT to study its functions in early embryos. Using this approach, we demonstrate that the slicer activity of CSR-1 is essential for embryonic viability and is required to post-transcriptionally cleaves its embryonic mRNA targets. Given the conservation of Argonaute catalytic activity in metazoans and the conserved function of small RNAs in regulating maternal mRNA clearance in several animal models we propose that a similar mechanism operates to clear maternal mRNAs during the MZT across species.

## RESULTS

### CSR-1 localizes to the cytoplasm of somatic blastomeres and its slicer activity is essential for embryonic development

To study CSR-1 localization during embryogenesis, we have generated various CRISPR-Cas9 tagged versions of CSR-1^23^. CSR-1 is expressed mainly in the germline of adult worms^17^ and localizes in the cytoplasm and germ granules (Extended Data Fig. 1a, b). However, in early embryos, CSR-1 localizes both in the germline and somatic blastomeres (Fig. 1a, Extended Data Fig. 1b). CSR-1 in the somatic blastomeres is mainly cytoplasmic and persists for several cell divisions (Fig. 1a, Extended Data Fig. 1b). This is in contrast with other germline Argonaute proteins^24,25^, such as PIWI, which exclusively segregate with the germline blastomere from the first embryonic cleavage (Fig. 1a, Extended Data Fig. 1b). Animals lacking CSR-1 or expressing a CSR-1 catalytic inactive protein show several germline defects, severely reduced brood sizes, chromosome segregation defects, and embryonic lethality^16,21,26^.

**Fig. 1:**
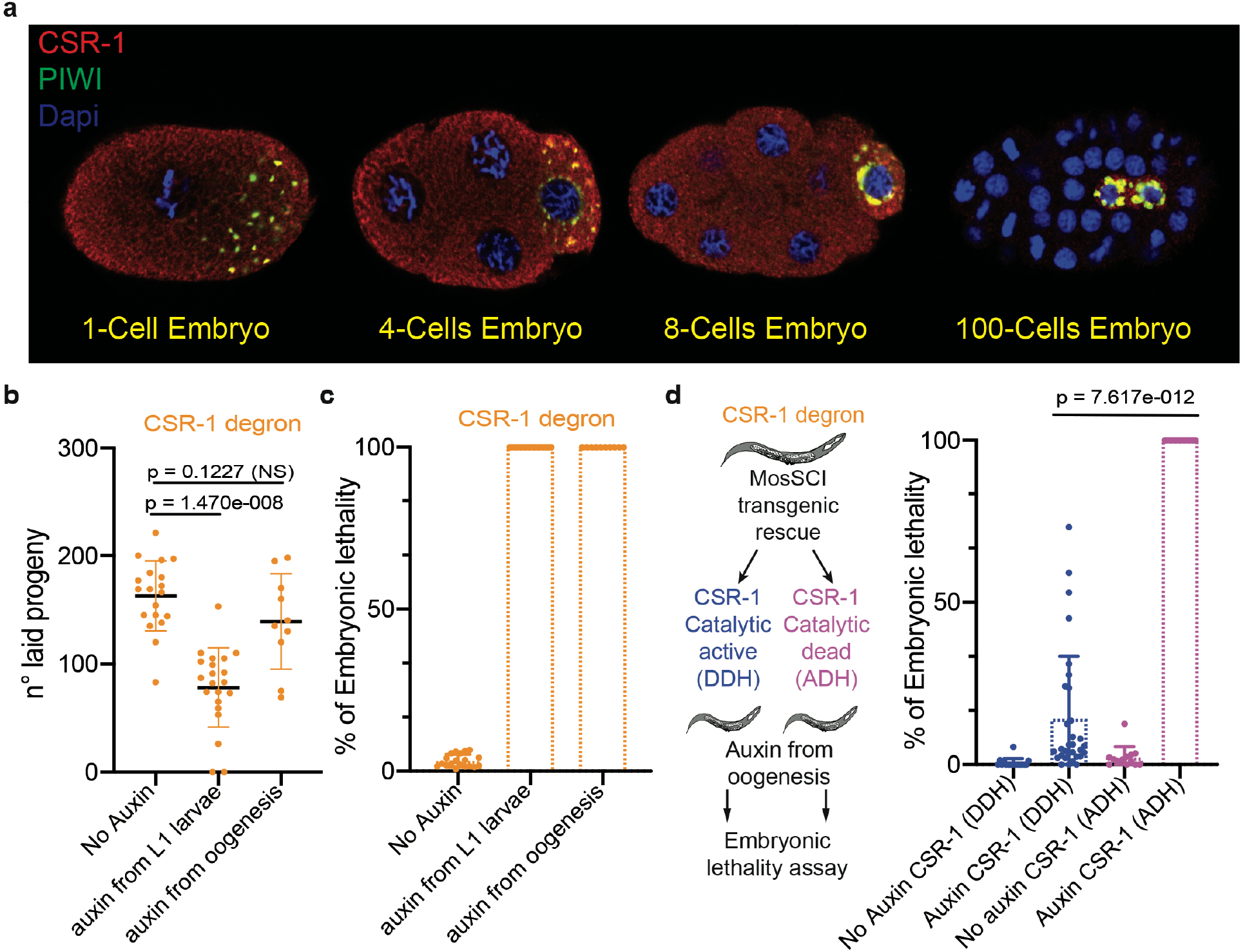
CSR-1 localizes to the cytoplasm of somatic blastomeres and its slicer activity is essential for embryonic development. **a**, Immunostaining of CSR-1 and PIWI in 1-cell, 4-cells, 8-cells, and more than 100-cells embryos of a 3xFLAG::HA::CSR-1 CRISPR-Cas9 strain, using anti-FLAG and anti-PRG-1 antibodies. DAPI signal is shown in blue. **b**, Brood size assays of CSR-1 degron strains with and without Auxin. The data points correspond to the number of living larvae from individual worms. Data are mean ± s.d. Two-tailed P values were calculated using Mann–Whitney–Wilcoxon tests. **c**, Percentages of embryonic lethality from the brood size experiment shown in **b**, measured as the percentage of dead embryos versus the total number among laid embryos. Data are mean ± s.d. **d**, Percentage of embryonic lethality in CSR-1 degron strains complemented with single-copy transgenic expression of CSR-1 with (ADH) or without (DDH) mutation in the catalytic domain. The percentage of embryonic lethality is calculated by dividing the number of dead embryos by the total number of laid embryos. Dots correspond to the percentages of embryonic lethality from individual worms. Data are mean ± s.d. Two-tailed P values were calculated using Mann–Whitney– Wilcoxon tests. The graphical representation of the experiment is shown on the left.

To understand the role of CSR-1 during embryogenesis, we depleted CSR-1 protein using the auxin-inducible degradation (AID) system^22^ (Extended Data Fig. 3a, b). First, we depleted CSR-1 from the first larval stage (L1), recapitulating the fertility loss and embryonic lethality phenotypes of both *csr-1 (tm892)* mutant and catalytically-inactive CSR-1 mutant worms^21,26^ (Fig. 1b, c, and Extended Data Fig. 2a, b). Next, we depleted CSR-1 from the beginning of oogenesis (Extended Data Fig. 3a, b). This treatment did not significantly decrease the fertility of adult worms (Fig. 1b), yet all the CSR-1-depleted embryos failed to develop in larvae (Fig. 1c), suggesting CSR-1 plays an essential role during embryogenesis.

To test whether the catalytically active CSR-1 is required for embryonic viability during the MZT, we expressed a single transgenic copy of CSR-1, using MosSCI^27^, with or without an alanine substitution of the first aspartate residues within the catalytic DDH motif and depleted endogenous CSR-1 at the beginning of oogenesis (Fig. 1d). The CSR-1 catalytic dead mutant (ADH) failed to suppress the embryonic lethality caused by the depletion of endogenous CSR-1 (Fig. 1d), whereas catalytically active CSR-1 (DDH) significantly restored embryonic viability (Fig. 1d). Thus, CSR-1 slicer activity is essential during embryogenesis.

### CSR-1 post-transcriptionally regulates its targets in the embryo

The localization of maternally inherited CSR-1 in the somatic blastomeres during early embryogenesis suggests that CSR-1 and its interacting 22G-RNAs may regulate specific embryonic targets in somatic cells. To better understand the regulatory function of CSR-1 in the embryo, we identified and compared CSR-1-interacting 22G-RNAs in the embryo and adult worms. CSR-1 was immunoprecipitated in embryo populations ranging from 1-to 20-cells stage (Extended Data Fig. 4a) or young adult worms, followed by sequencing of interacting 22G-RNAs. CSR-1-22G-RNA targets in the adult germlines and early embryos largely overlap (74 %) (Fig. 2a). However, the relative abundance of the 22G-RNAs complementary to these targets was different between the adult germlines and early embryos (Fig. 2a, b). Only 29 % of CSR-1 targets with the highest levels of complementary 22G-RNAs (> 150 reads per million, RPM) in adults and embryos are shared (Fig. 2a, b). Given that CSR-1 cleavage activity on target mRNAs depends on the abundance of 22G-RNAs^21^, our results suggest that CSR-1 may regulate a different subset of mRNAs in the embryos.

**Fig. 2:**
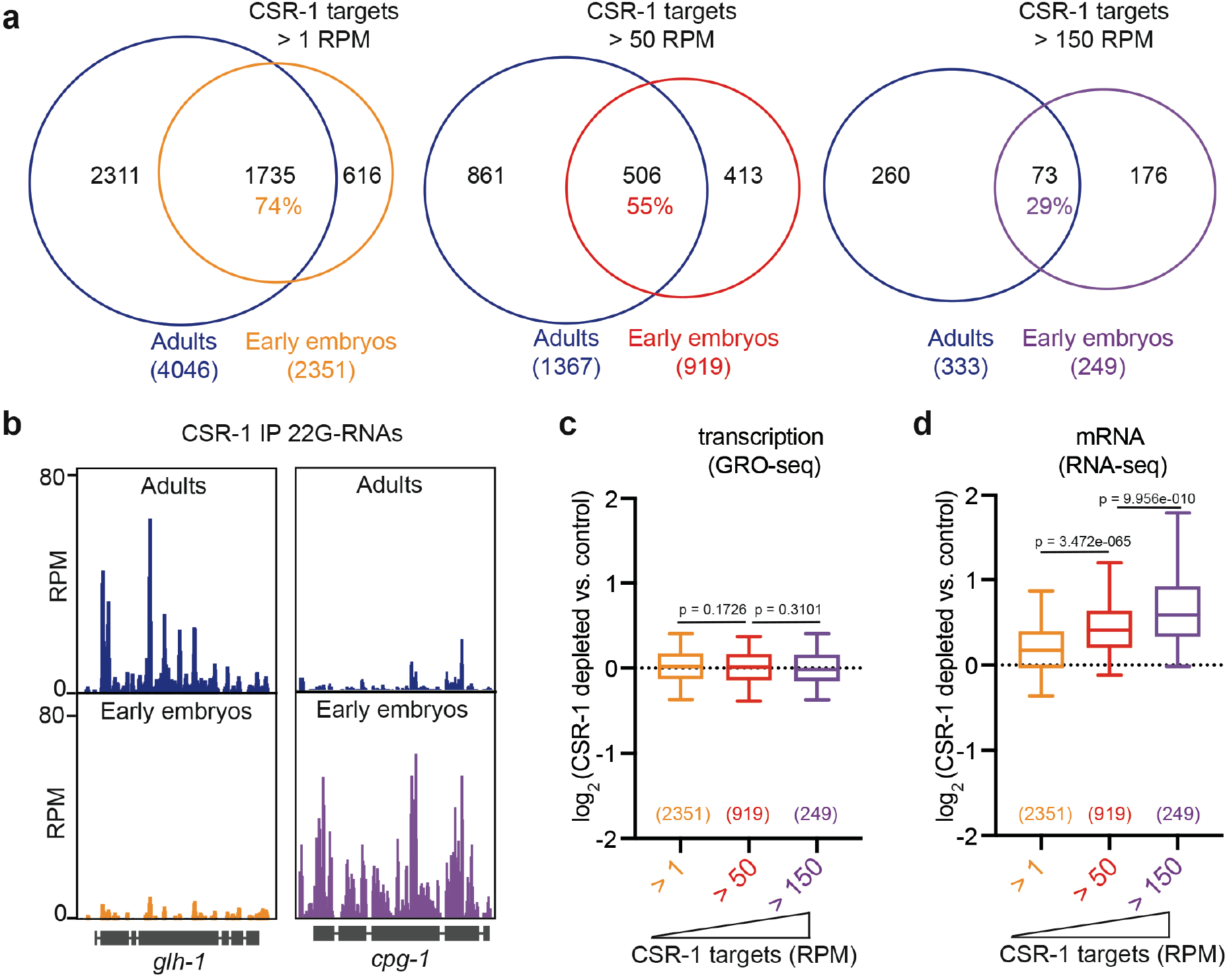
CSR-1 post-transcriptionally regulates its targets in the embryo. **a**, Venn diagram showing the overlap between CSR-1 targets in adults and in early embryos. A gene is considered a CSR-1 targets when the ratio between normalized 22G-RNAs from CSR-1 immunoprecipitation and the total RNA in the input is at least 2. We have selected three categories of CSR-1 targets based on the density (reads per million, RPM) of 22G-RNAs: CSR-1 targets > 1 RPM, CSR-1 targets > 50 RPM, CSR-1 targets > 150 RPM. The number of genes and the percentage of overlapping is indicated. **b**, Genomic view of two CSR-1 target genes showing normalized 22G-RNAs in CSR-1 IPs from adult (top) or early embryos (bottom). **c**, **d**, Box plots showing the log2 fold change of nascent RNAs (by GRO-seq) or mRNAs (by RNA-seq) in CSR-1 depleted early embryos compared to control. The distribution for the embryonic CSR-1 targets > 1 RPM, CSR-1 targets > 50 RPM, CSR-1 targets > 150 RPM are shown. Box plots display median (line), first and third quartiles (box) and highest/lowest value (whiskers). Two-tailed P values were calculated using Mann–Whitney–Wilcoxon tests.

To evaluate gene expression changes upon CSR-1 depletion in embryos, we depleted CSR-1 from the beginning of oogenesis and collected early embryo populations fully depleted of CSR-1 (Extended Data Fig. 3a, and Extended Data Fig. 4b, c). Analysis of nascent transcription using global run-on sequencing (GRO-seq) revealed that the zygotic transcription in CSR-1 depleted embryos was largely unaffected compared to control untreated embryos (Fig. 2c, Extended Data Fig. 4b), indicating that CSR-1 depleted embryos could initiate zygotic transcription. However, analysis of steady-state mRNA levels using RNA-seq revealed increased levels of CSR-1 embryonic mRNA targets (Fig. 2d, Extended Data Fig. 4c), which correlated with the levels of complementary antisense 22G-RNAs loaded into CSR-1 (Fig. 2d). These results suggest that inherited CSR-1 22G-RNAs in the embryo, targets a different set of mRNAs compared to adult germlines and these new target mRNAs are directly regulated at the post-transcriptional level in a dose-dependent manner.

### CSR-1 cleaves maternal mRNAs in early embryos

In the first phases of embryogenesis, the embryo is transcriptionally silent and relies on maternally inherited mRNA transcripts. Maternal mRNAs degradation is concomitant with the zygotic genome activation, during the MZT^1^. To gain insight into the type of mRNA targets regulated by CSR-1, we asked whether CSR-1 embryonic targets corresponded to maternal or zygotic mRNA transcripts. We adapted a previously developed sorting strategy^28^ to sort embryo populations at the 1-cell stage (99 % pure), or enrich for early embryos (from 4- to 20-cells), or late embryos (more than 20-cells) (Extended Data Fig. 4d). Strand-specific RNA-seq was performed on these embryo populations to profile gene expression dynamics. We detected mRNAs from 7033 genes in 1-cell embryos (transcript per million, TPM, > 1) (Supplemental Table 1), which we classified into three groups based on their dynamics during embryonic development. “Maternal cleared” mRNAs (1320 genes) corresponded to maternal mRNAs in 1-cell embryos whose levels diminished in early and late stages (Fig. 3a) (Supplemental Table 1). These genes belong to the previously characterized Maternal class II genes, which are maternally inherited in the embryos and degraded in somatic blastomeres^12^. “Maternal stable” mRNAs (1020 genes) corresponded to maternal mRNAs in 1-cell embryos whose levels are stably maintained in early and late embryos (Fig. 3a) (Supplemental Table 1). “Zygotic” mRNAs (704 genes) whose mRNAs accumulated in early and late embryos but were undetectable in 1-cell embryos (TPM < 1) (Fig. 3a) (Supplemental Table 1).

**Fig. 3:**
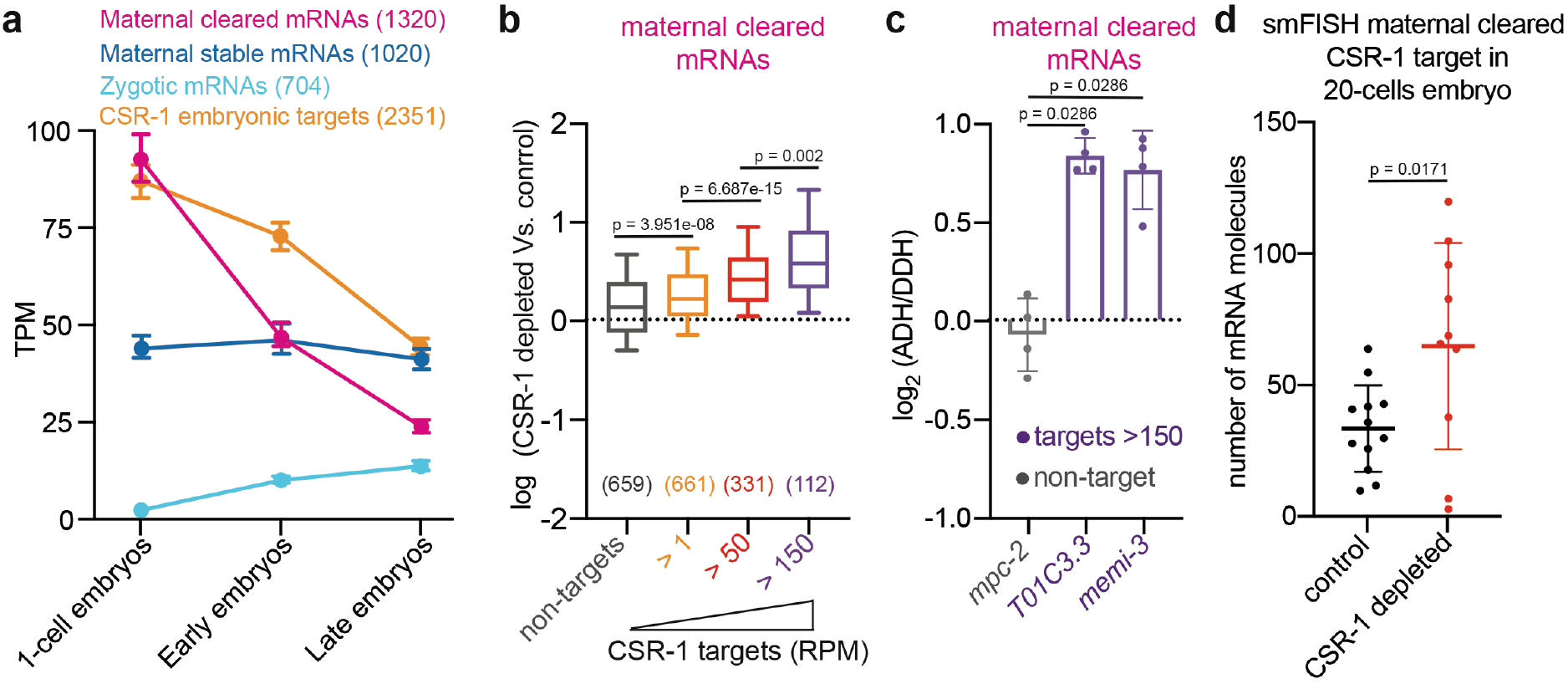
CSR-1 cleaves maternal mRNAs in early embryos. **a**, Detection of maternal cleared mRNAs, maternal stable mRNAs, zygotic mRNAs, and CSR-1 embryonic target mRNAs by RNA-seq from sorted 1-cell stage, early stage, and late stage embryos. Median levels and 95% confident interval of normalized read abundances in transcript per million (TPM) are shown. **b**, Box plots showing the log2 fold change of cleared maternal mRNAs in CSR-1 depleted early embryos compared to control. The distribution for the non-targets, CSR-1 targets > 1 RPM, CSR-1 targets > 50 RPM, CSR-1 targets > 150 RPM are shown. Box plots display median (line), first and third quartiles (box) and highest/lowest value (whiskers). **c**, RT–qPCR showing log2 fold change of maternal cleared mRNAs in CSR-1 depleted embryos with transgenic expression of CSR-1 with (ADH) or without (DDH) mutation in the catalytic domain. Data are mean ± s.d. and the dots individual data from four biologically independent experiments. **d**, smFISH quantification of *C01G8.1* mRNA target in somatic blastomeres of 20-cell stage embryos depleted of CSR-1 compared to the control. Data are mean ± s.d. and the dots number of mRNA molecules per embryo section. In all figures two-tailed P values were calculated using Mann– Whitney–Wilcoxon tests.

Analysis of the embryonic CSR-1 targets revealed that they were inherited in 1-cell embryos and gradually depleted in early and late embryos, similarly to the maternal cleared class of genes (Fig. 3a), suggesting that CSR-1 contributes to maternal mRNA clearance in somatic blastomeres. To validate this result, we analyzed the levels of maternal cleared mRNAs in CSR-1 depleted embryos and found accumulation of maternal cleared mRNA targets correlated with the abundance of antisense 22G-RNAs loaded onto CSR-1 (Fig. 3b). To determine if CSR-1 slicer activity is required to regulate those targets we measured gene expression changes in CSR-1 depleted embryos complemented with transgenic expression of CSR-1 catalytic dead mutant (ADH) or catalytically active CSR-1 (DDH), and found increased accumulation of CSR-1 targets in slicer-inactive CSR-1 embryo population (Fig. 3c). Finally, we performed RNA-seq on CSR-1 depleted in 1- to 4-cells stage enriched embryos or 4- to 20 cell stage enriched embryos (Extended Data Fig. 5a-c) to study if CSR-1 is directly regulating these maternal mRNA targets in embryos. CSR-1 depletion in 1- to 4-cell embryos did not cause global changes in maternal cleared mRNA target levels (Extended Data Fig. 5d). However, CSR-1 depletion in 4- to 20-cell stage enriched embryos resulted in increased levels of maternal cleared mRNA targets compared to 1- to 4-cell embryos (Extended Data Fig. 5d). Thus, CSR-1 and its interacting 22G-RNAs actively degrade their maternal mRNA targets during early embryogenesis. To verify these results, we performed RNA single-molecule fluorescence *in situ* hybridization (smFISH) on a maternal cleared CSR-1 mRNA target in CSR-1 depleted and control embryos. We observed reduced maternal mRNA degradation of the CSR-1 target in somatic blastomeres of 20-cell stage embryos depleted of CSR-1 compared to control (Fig. 3d, Extended Data Fig. 5e). Collectively, these results suggest that the maternally inherited CSR-1 loaded with 22G-RNAs contribute to cleave and degrade complementary maternal mRNA targets in somatic blastomeres.

### CSR-1 preferentially targets maternal mRNAs no longer engaged in translation

To analyze the dynamics of CSR-1-dependent mRNA clearance during embryogenesis, we divided maternally inherited mRNAs into those degraded early and late during embryogenesis. Early-degraded mRNAs (483 genes) showed at least a 2-fold reduction in mRNA levels in sorted early embryo populations compared to 1-cell embryos (Fig. 4a, Extended Data Fig. 4d). Late-degraded mRNAs (1572 genes) showed stable levels of mRNAs in sorted early embryo populations compared to 1-cell embryos but were decreased at least 2-fold in populations of late embryos (Fig. 4a, Extended Data Fig. 4d). Analysis of 22G-RNAs interacting with CSR-1 in the embryo showed that the levels of 22G-RNAs targeting the early-degraded mRNAs were significantly higher compared to late-degraded mRNAs in total RNAs and CSR-1 IPs (Extended Data Fig. 6a, b). These results suggest that the abundance of 22G-RNAs correlates with the temporal decay of maternal cleared mRNAs during embryogenesis.

**Fig. 4:**
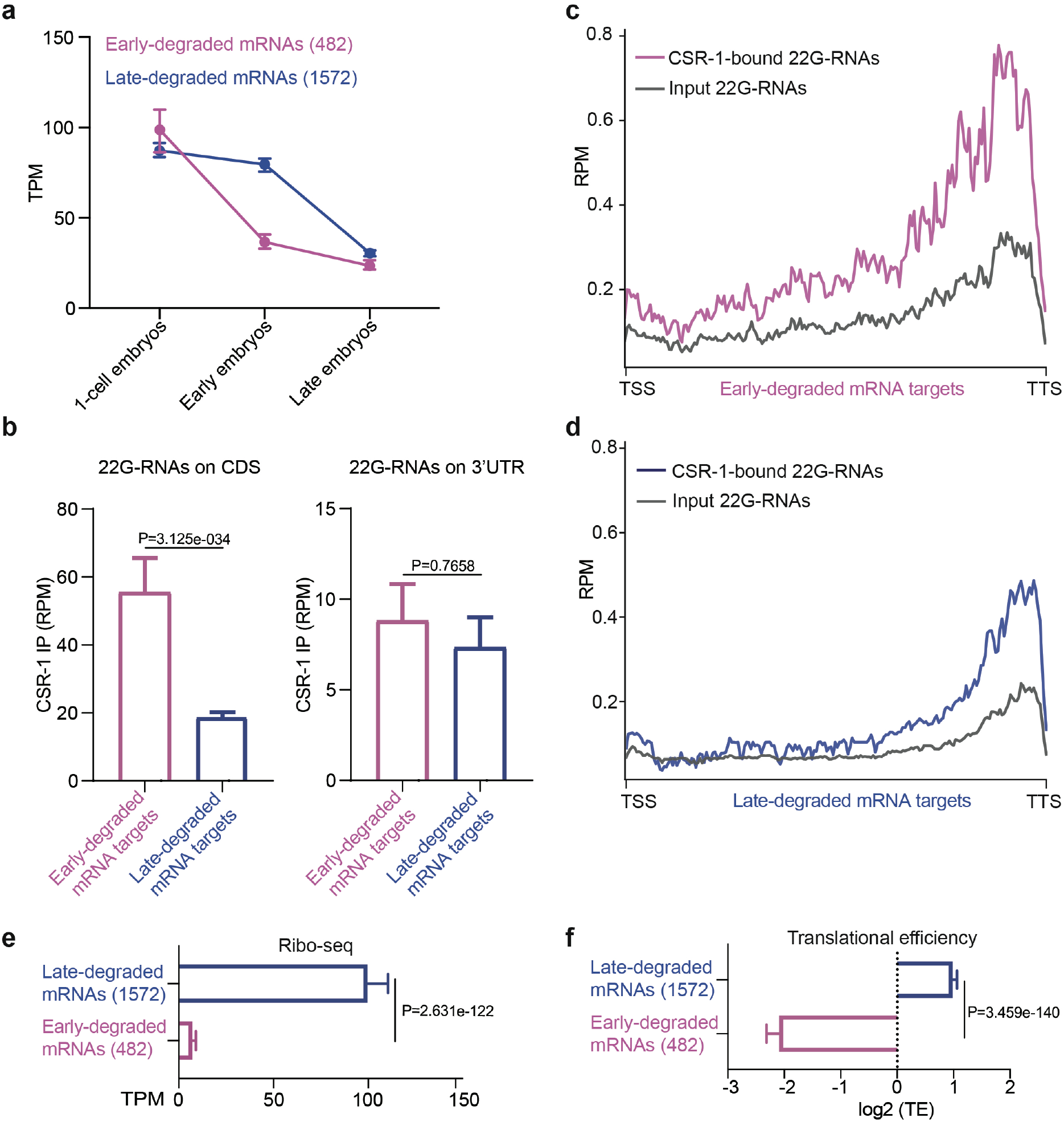
CSR-1 preferentially targets maternal mRNAs no longer engaged in translation. **a**, Detection of early- and late-degraded mRNAs by RNA-seq from sorted 1-cell, early and late embryo stages. Median levels and 95 % confident interval (CI) of normalized read abundances in transcript per million (TPM) are shown. **b**, Abundance of CSR-1-bound 22G-RNAs antisense to coding sequence or 3’UTR of early and late-degraded mRNAs. Median levels and 95 % CI of normalized showing normalized 22G-RNA reads (RPM) are shown. **c**, **d**, Metaprofile analysis showing normalized 22G-RNA reads (RPM) across early-(magenta) and late-degraded (blue) mRNA targets in CSR-1 immunoprecipitation (IP) or total RNA input. The transcriptional start site (TSS) and transcriptional termination site (TTS) is indicated. **e**, Ribosomal occupancy (by Ribo-seq) of early- and late-degraded mRNAs in early embryos. Median levels and 95 % CI of normalized ribosomal protected fragment (RPF) are shown. **f**, Median levels and 95 % CI of translational efficiency (TE) of early- and late-degraded mRNAs in early embryos. Two-tailed p value in **d, e**, were calculated using the Mann-Whitney-Wilcoxon test.

We generated metaprofiles for the levels of CSR-1-interacting 22G-RNAs levels along the gene body of early- and late-degraded mRNA targets (Fig. 4c, d). The early-degraded mRNA genes showed an enrichment of CSR-1-loaded 22G-RNAs along the whole coding sequence (Fig. 4b, c). The late-degraded mRNA genes showed enrichment of CSR-1-loaded 22G-RNAs mainly at the 3’ end, corresponding to the 3’UTR (Fig. 4b, d). Previous reports have shown that the rate of mRNA clearance during maternal to zygotic transition is influenced by translational efficiency^29,30^. Thus, the different translational status of early- and late-degraded mRNAs may influence the accessibility and production of CSR-1-loaded 22G-RNAs antisense to the coding sequences of cleared mRNAs. To test this hypothesis, we measured ribosomal occupancy by Ribo-seq in a population of early embryos (Extended Data Fig. 6c). Early-degraded mRNAs showed significantly lower ribosomal occupancy and translational efficiency compared to late-degraded mRNAs (Fig. 4e, f). These results suggest that ribosomal loading on maternal mRNAs can influence their coding sequence accessibility for the 22G-RNA-mediated cleavage by CSR-1 in somatic blastomeres.

### Ribosome occupancy affects CSR-1-mediated maternal mRNA clearance

Previous studies have shown that mRNA codon usage can influence translational efficiency and maternal mRNA decay during the MZT^29,30^. However, the early- and late-degraded mRNAs did not show any differences in their codon usage (Extended Data Fig. 6d, e). Because the translation of germline mRNAs is primarily regulated by their 3’UTR in *C. elegans*^31^, we investigated whether different 3’UTRs from early- and late-degraded mRNAs can impact translation and mRNA degradation in early embryos. We generated two single-copy transgenic lines expressing a germline *mCherry::h2b* mRNA reporter with either a 3’UTR from an early-(*egg-6* 3’UTR) or a late-(*tbb-2* 3’UTR) degraded mRNA target (Fig. 5a). The steady-state level of the two *mCherry::h2b* mRNA reporters was similar in adult worms (Fig. 5b), whereas the level of *mCherry::h2b* mRNA fused with the early-degraded *egg-6* 3’UTR was lower in embryo, indicating increased degradation in early embryos (Fig. 5b). To verify that the different 3’UTRs affect the translational efficiency of *mCherry::h2b* mRNA in embryos we quantified the ribosome occupancy by Ribo-seq and mRNA level by RNA-seq, and observed decreased translational efficiency of *mCherry* transgenic reporter with *egg-6* 3’UTR (Fig. 5c, Extended Data Figs. 6f and 7a, b). To test whether CSR-1 22G-RNAs contribute to the increased decay of the *mCherry::h2b* reporter fused to the *egg-6* 3’UTR, we sequenced small RNAs in early embryo preparations from the two transgenic lines (Extended Data Fig. 6f). We observed increased levels of small RNAs on the coding sequences of the *mCherry* transgenic reporter with *egg-6* 3’UTR compared to the *tbb-2* 3’UTR reporter (Fig. 5d, e). Thus, the translation level of maternal mRNAs can influence the rate of CSR-1-mediated mRNA cleavage in somatic blastomeres.

**Fig. 5:**
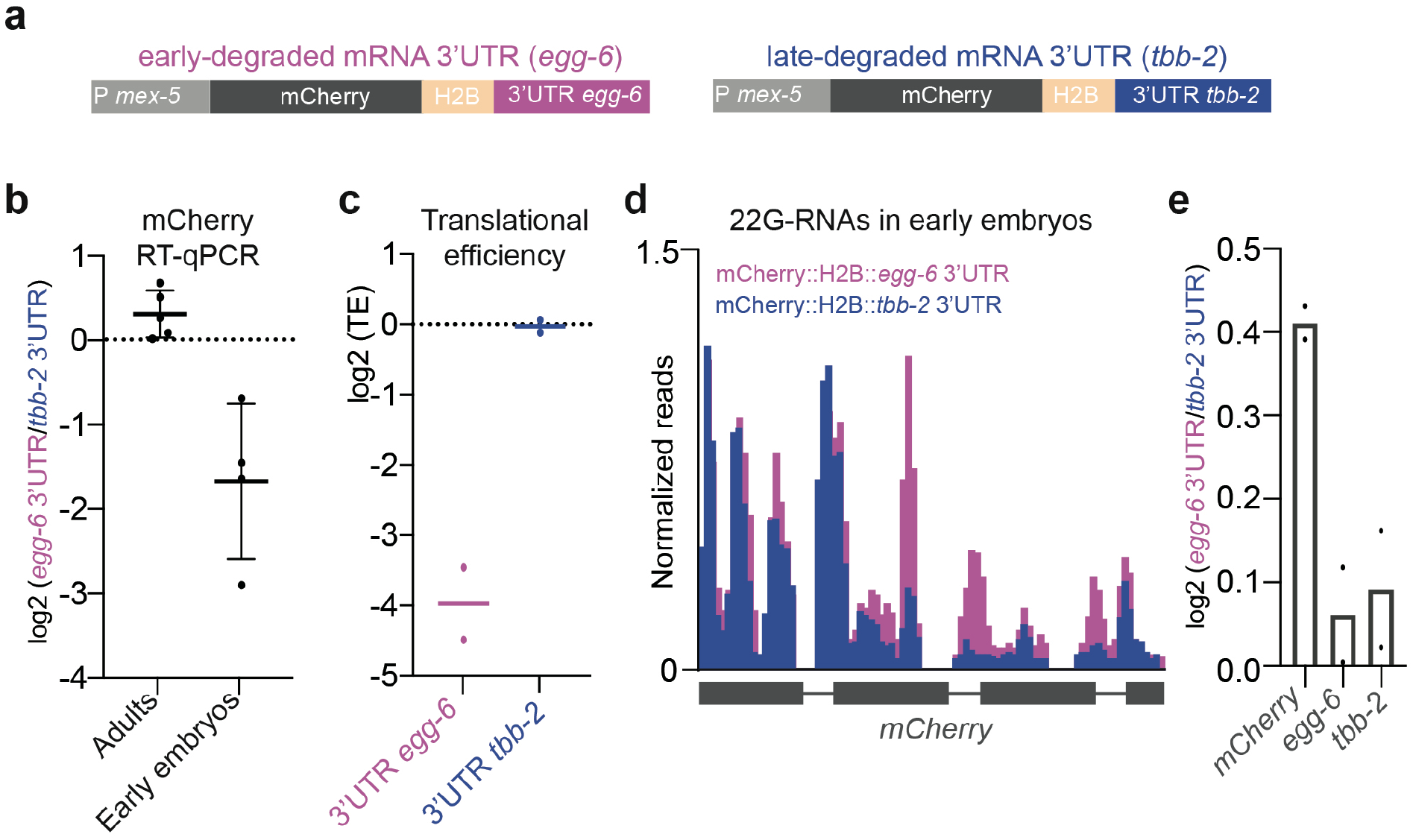
Ribosome occupancy affects CSR-1-mediated maternal mRNA clearance. **a**, Schematic of the germline-expressed single-copy *mCherry::h2b* transgenic mRNA fused to 3’UTR derived from early- (*egg-6 3’UTR*) or late-degraded (*tbb-2 3’UTR*) mRNAs. **b**, RT-qPCR assay to detect mRNAs from the two transgenic reporters in adults and early embryos. The black lines indicate the mean, the error bars the standard deviation, and the black dots individual data from four biologically independent experiments. **c**, Translational efficiency (TE) of *mCherry::h2b* transgenic mRNAs. The lines indicate the mean and the dots individual data from two biologically independent experiments. **d**, Genomic view of normalized 22G-RNAs antisense to *mCherry*. **e**, 22G-RNA log2 fold change antisense to *mCherry*, *egg-6* and *tbb-2* genes in *egg-6 3’UTR* vs. *tbb-2 3’UTR* transgenes. The bars indicate the mean and the dots individual data from two biologically independent experiments.

In summary, our study identifies a novel function of the conserved Argonaute slicer activity in the degradation of maternally inherited mRNAs during the MZT.

## DISCUSSION

In this study, we have shown that the maternally inherited CSR-1 protein and its interacting 22G-RNAs trigger the cleavage of hundreds of complementary maternal mRNA targets in embryos. Despite most of the inherited Argonautes localize to the germline blastomeres, the inherited CSR-1 also localizes to the cytoplasm of somatic blastomeres for several cell divisions during early embryogenesis. We have shown that *C. elegans* embryos inherit a pool of 22G RNAs, loaded onto CSR-1, antisense to mRNAs that undergoes rapid degradation in developing embryos, and CSR-1 slicer activity facilitates the clearance of these maternal mRNAs in somatic blastomeres. We have observed that CSR-1 is preferentially loaded with 22G-RNAs antisense to the coding sequences of untranslated mRNA targets in embryos. Based on our results, we propose that CSR-1 slicer activity is required during embryogenesis to cleave untranslated maternal mRNAs accumulated in somatic blastomeres after the first embryonic divisions. We speculate that active translation prevents the accessibility of CSR-1 22G-RNAs on complementary mRNA targets and hence the slicing through the coding sequences of translated maternal mRNAs (Fig. 6). Therefore, the translation of maternal mRNAs ensures that CSR-1 22G-RNA complexes can only remove these mRNAs in the somatic cells of the developing embryos, when they are not needed to be translated anymore. This antagonism between translation and CSR-1 cleavage might explain why CSR-1 slicer activity results in only fine-tuning some of its complementary targets in the adult germlines. Thus, the presence of ribosomes on germline mRNAs may prevent the slicer activity of CSR-1, which is instead permitted in the embryos after the mRNAs are disengaged from translation and can be degraded. We have also shown that by modulating the translational activity of a maternal mRNA sequence we could detect different amounts of antisense 22G-RNAs. Therefore, it is possible that the presence of ribosome on the coding sequence of maternal mRNAs can antagonize the synthesis of 22G-RNAs by the RdRP.

**Fig. 6:**
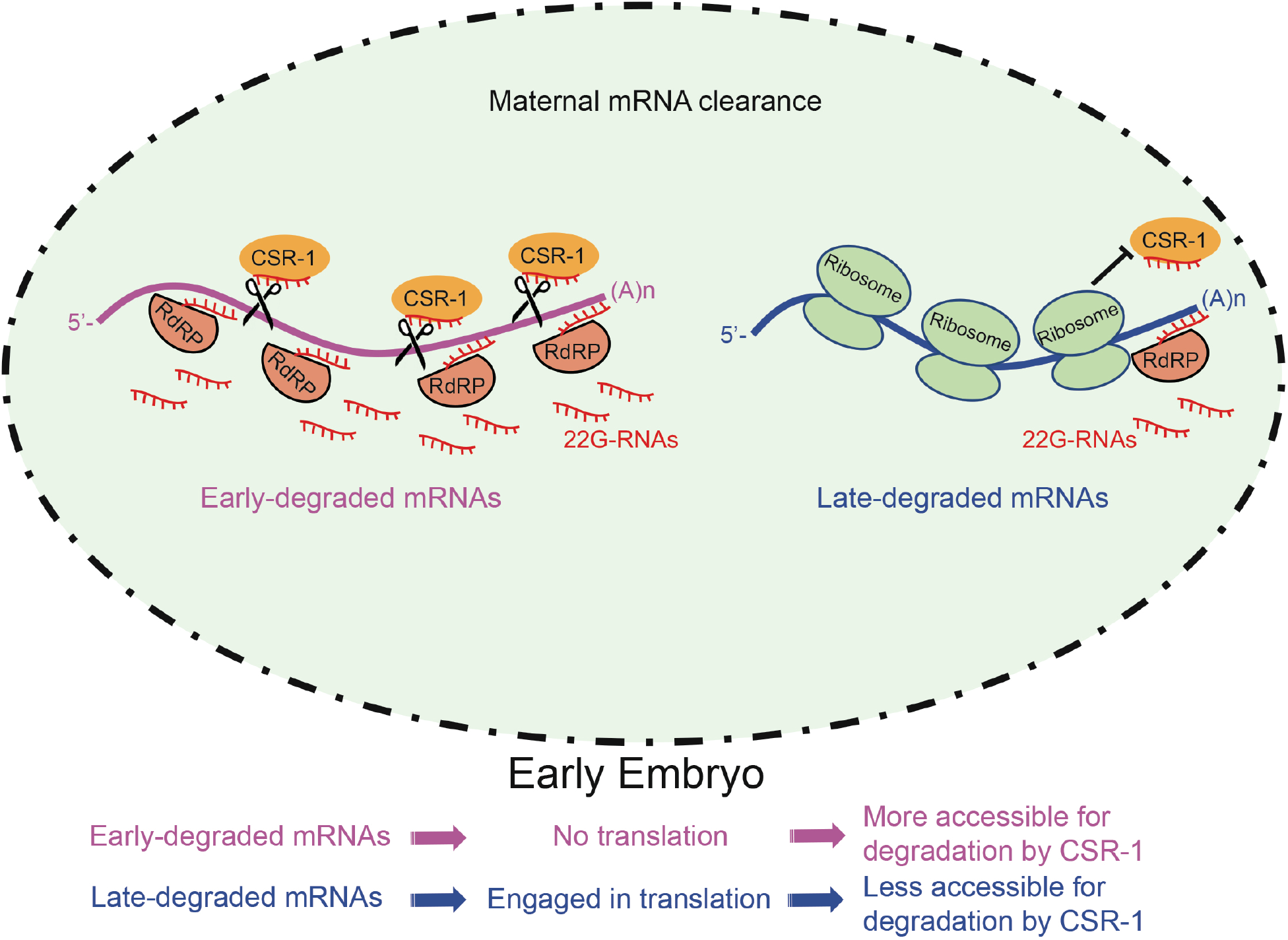
Model illustrating CSR-1-dependent clearance of maternal mRNA in *C. elegans* embryos.

It is known that translation affects maternal mRNA clearance, and recent works have shown how the codon usage of maternal mRNAs regulates translation and degradation rates in zebrafish embryos^29,30^. Here we show that in *C. elegans* embryos early and late degraded maternal mRNAs do not differ in their codon usage. Instead, we have observed that the 3’UTR is sufficient to regulate the translational efficiency of the coding sequence of a maternal mRNA in embryos, and in turn its degradation rate, including CSR-1 slicer activity. A consensus sequence in the 3’UTR of some maternal mRNAs has been shown to play a role in the degradation of mRNAs from oocytes to 1-cell embryos in *C. elegans*^9^. Therefore, it would be interesting in the future to identify other consensus sequences located in the 3’UTR of maternal mRNAs and also RNA binding proteins that might contribute together with CSR-1 to regulate mRNA clearance during embryogenesis.

CSR-1 has been extensively studied in the adult germlines, where it has been proposed to protect active genes from piRNA silencing^18,19^, promote germline transcription^32,33^, regulate the biogenesis of histone mRNAs^34^, and fine-tune germline mRNAs loaded into oocytes^21^. The catalytic activity of CSR-1 and its gene regulatory function on target mRNAs has been proven be essential for fertility and chromosome segregations^21^. Similarly, our work provided evidences that CSR-1 slicer activity is essential for embryonic viability independently from its germline functions. Therefore, we propose that the accumulation of maternal mRNAs in embryos lacking CSR-1 slicer activity might be deleterious for the developing embryo. Moreover, our finding suggests that mRNA translation in the germline might inhibit CSR-1 slicer activity on target mRNAs and results only in a mild fine-tuning of some targets. Therefore, we propose that the role of CSR-1 in protecting germline mRNA from piRNA silencing might be still compatible with its cleavage activity.

The requirement of Argonaute slicer activity to degrade endogenous target mRNAs has only been described in the silencing of repetitive elements in animals. However, catalytically active Argonaute proteins are conserved in animals, including humans. We propose that the RNAi activity of CSR-1 on maternal mRNAs might be a conserved mechanism required for the MZT and mRNA clearance across species. Even if miRNAs contribute to regulate maternal mRNA clearance in some animal models, including *Drosophila*, zebrafish and *Xenopus*, their functions are suppressed in mouse oocytes^7,8^. Instead, endogenous small RNAs targeting protein-coding genes, similar to the one loaded onto CSR-1, have been detected in mouse oocytes and embryonic stem cells^35–37^. This suggests that mammalian RNAi, in addition to roles in the suppression of repetitive elements, might also regulate endogenous genes. Given that the inheritance of maternal Ago2, the only catalytically active Argonaute in mouse, is essential for oocyte^10^ and early embryonic development^11^, the current hypothesis is that Ago2 can promote the degradation of maternal mRNAs in embryos (Svoboda and Flemr, 2010). Strikingly, catalytically inactive Ago2 in mouse oocytes causes chromosome segregation and spindle assemble defects^10^ similar to CSR-1 catalytic mutant^21^, suggesting they might regulate common functional mRNA targets during oogenesis. Therefore, Ago2 and endogenous small RNAs might play a role in maternal mRNA clearance during the MZT in mouse.

Based on our findings, we propose that endogenous small RNAs and Argonaute-mediated cleavage of mRNA may be a conserved mechanism in metazoans to degrade maternal mRNAs in oocyte and embryos.

## ACKNOWLEDGEMENTS

We would like to thank all the members of the Cecere laboratory and Nicola Iovino for the helpful discussions on the paper and suggestions on the project, and G. Riddihough (Life Science Editors) for editing the manuscript. We thank the Mello and the Dumont laboratories for sharing strains and reagents. Some strains were provided by the CGC, funded by NIH Office of Research Infrastructure Programs (P40 OD010440). This project has received funding from the Institut Pasteur, the CNRS, and the European Research Council (ERC) under the European Union's Horizon 2020 research and innovation programme under grant agreement No ERC-StG- 679243. E.C. was supported by a Pasteur-Roux Fellowship program. P.Q. was supported by Ligue Nationale Contre le Cancer (S-FB19032).

## AUTHOR CONTRIBUTIONS

G.C. identified and developed the core questions addressed in the project with the contribution of P.Q., and wrote the paper with the contribution of P.Q., E.C. and M.S. P.Q. performed most of the experiments and analyzed the results together with G.C. E.C. and L.B. generated all the lines used in this study with the help of P.Q. M.S. performed the immunoprecipitation of CSR-1 and small RNA sequencing in adult worms and performed the ribo-seq experiments together with P.Q. B.L. performed all the bioinformatic analysis. F.M. analyzed the smFISH experiments. G.C. C.D. helped P.Q. in setting up the embryo FACS sorting.

## DATA AND MATERIALS AVAILABILITY

All the sequencing data are available at the following accession numbers GSE146062 (secure token for reviewers: krkpoycihjulhax). All other data supporting the findings of this study are available from the corresponding author on reasonable request.

---

**Extended Data Fig. 1:**
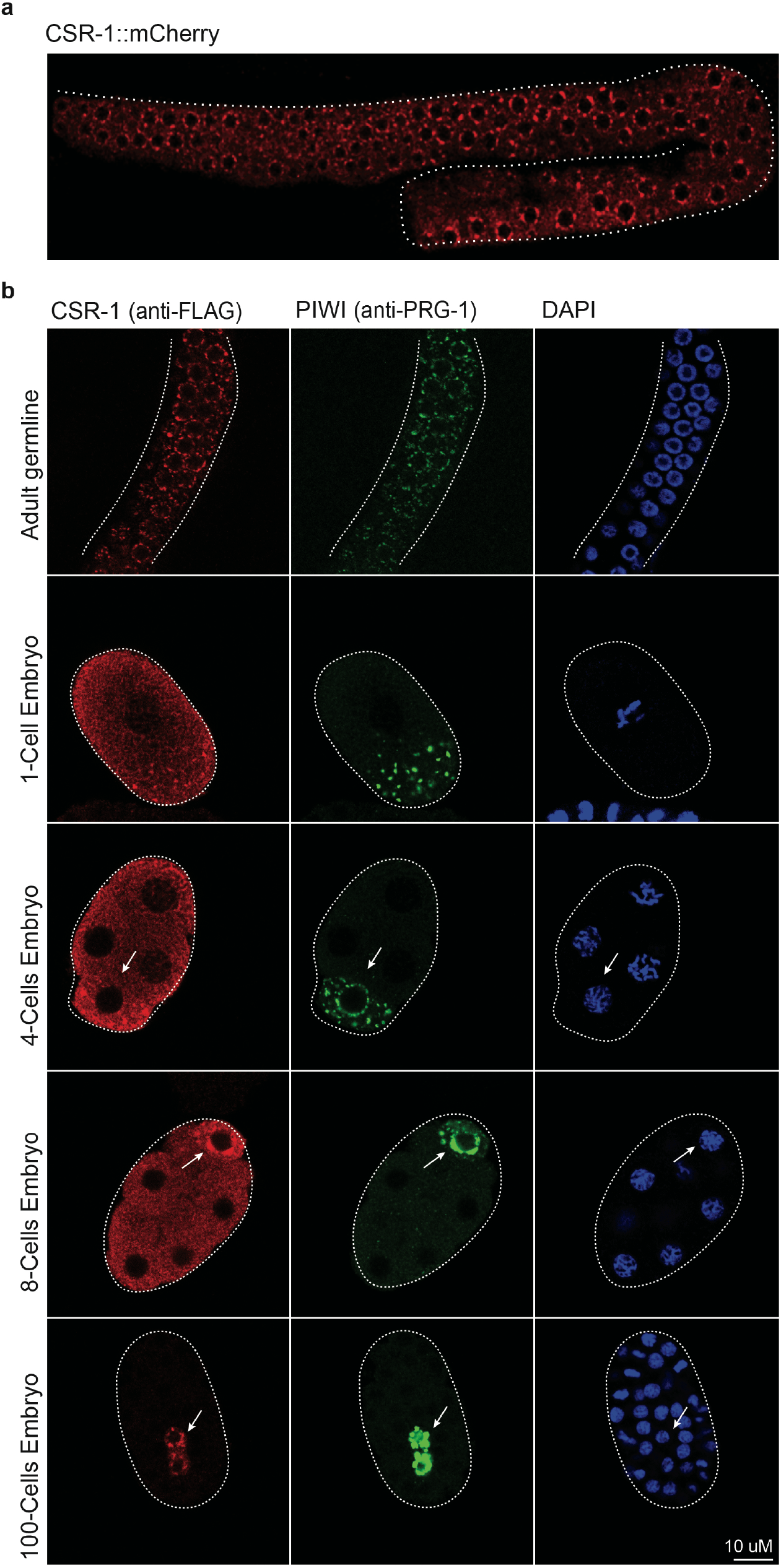
CSR-1 and PIWI localization in adult germlines and early embryos. **a**, Confocal image of CSR-1::mCherry in a gonad of late L4 larvae stage. (B) Immunostaining of CSR-1 and PIWI in dissected Adult gonad (top images) and 1-cell, 4-cells, 8 cells, and more than 100-cells embryo (bottom images) as shown in **Fig. 1a** using anti-FLAG and anti-PRG-1 antibodies. DAPI signal is shown in blue. Scale bar represent 10 μm. Arrows indicate germline blastomeres.

**Extended Data Fig. 2:**
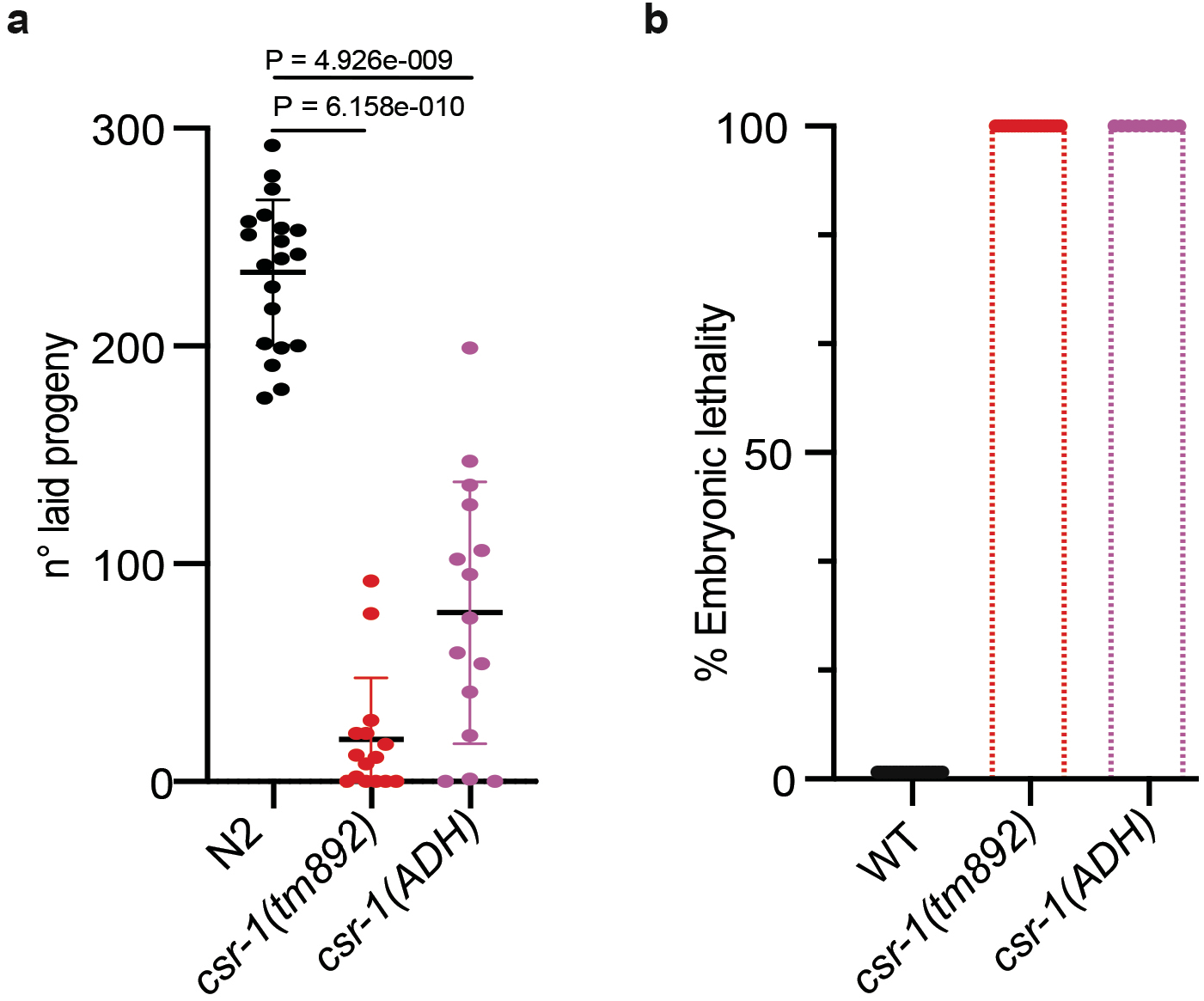
*csr-1(tm892)* mutant and CSR-1 catalytic dead (ADH) show fertility defects and embryonic lethality. **a**, Brood size assays of *csr-1(tm892)* mutant, CSR-1 catalytic dead (ADH) and wild-type. The data points correspond to the number of living larvae from individual worms. Data are mean ± s.d. Two-tailed P values were calculated using Mann–Whitney–Wilcoxon tests. **b**, Percentage of embryonic lethality from the brood size experiment shown in **a**, measured as the percentage of dead embryos versus the total number of laid embryos.

**Extended Data Fig. 3:**
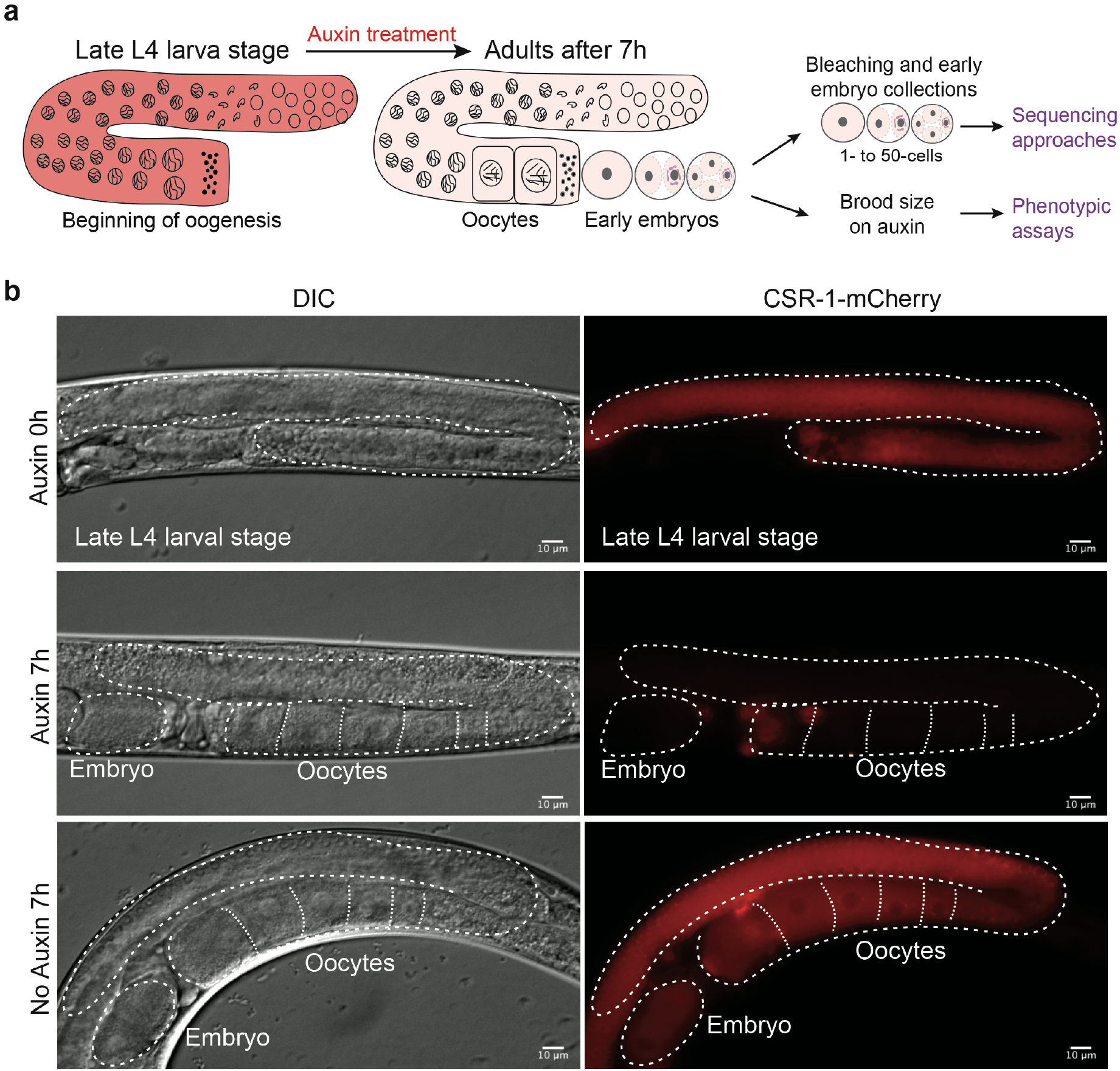
Depletion of CSR-1 during the MZT using the auxin-inducible degradation (AID) system. **a**, Schematic of auxin depletion strategy to obtain early embryo populations fully depleted of CSR-1. **b**, DIC micrographs (left) and fluorescent micrographs (right) of live animals expressing Degron::mCherry::3xFLAG::HA::CSR-1 used for auxin depletion experiments (see Materials and methods). Top images show animals before the treatment. Bottom images show gravid animals after treatment with ethanol (no auxin, control) or auxin. Auxin treatment completely depletes CSR-1 in adult gonads and embryos. Dashed lines represent gonads, oocytes and embryos. All scale bars represent 10 μm.

**Extended Data Fig. 4:**
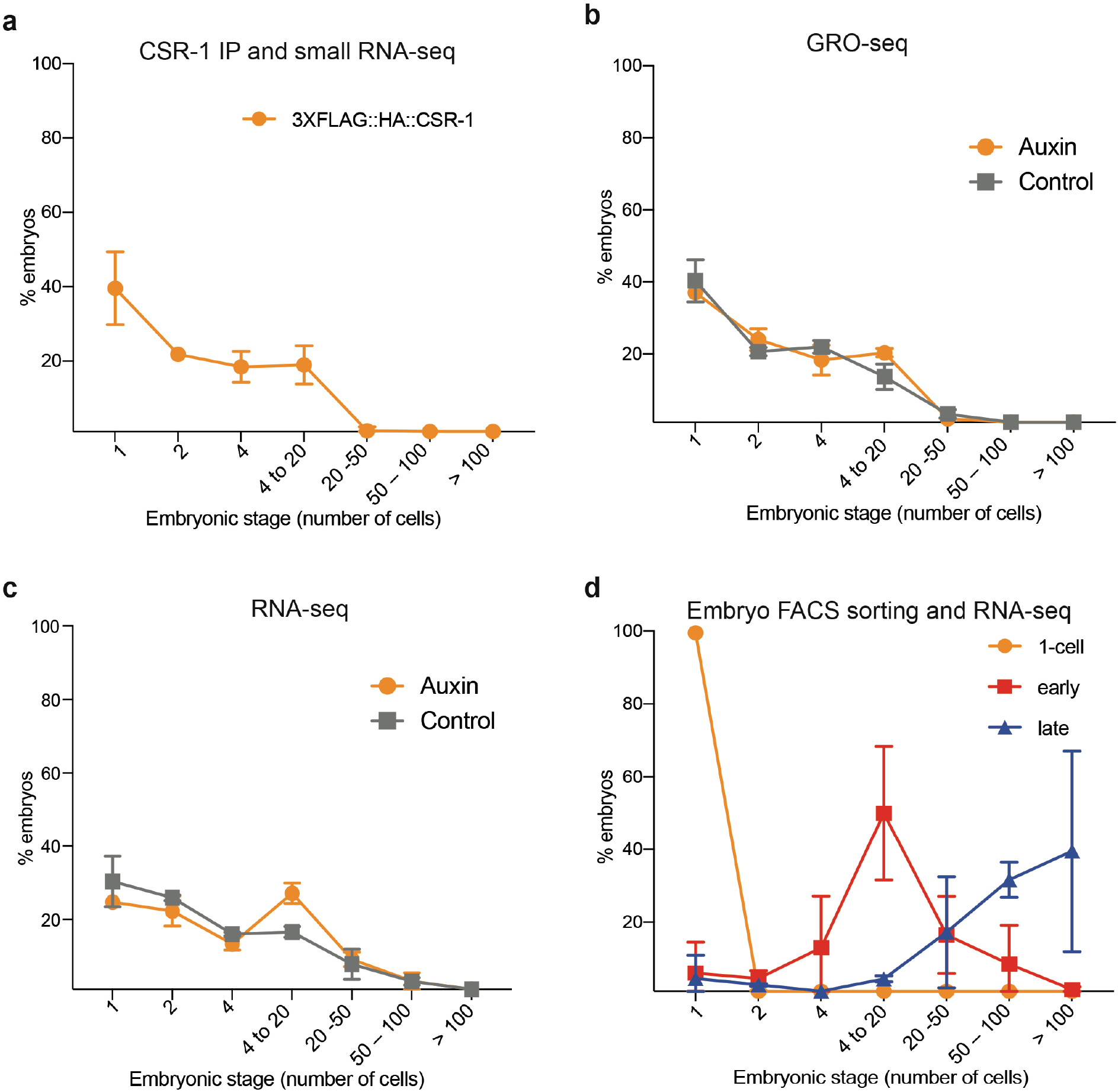
Early embryo populations used for sequencing experiments. **a**-**d**, Composition of embryo populations used for sequencing experiments. Embryos were stained with DAPI, frozen to stop cell divisions, and counted using a fluorescent microscope (see Material and Methods). **a**, 3xFLAG::HA::CSR-1 embryos used for small RNA-seq in **Fig. 2a, b**. **b, c,** Degron::mCherry::3xFLAG::HA::CSR-1 embryos treated with auxin or ethanol (No Auxin) used for GRO-seq **b**, and RNA-seq **c**, in **Fig. 2c, d,** and **Fig. 3b**. **d**, Embryos expressing OMA-1::mCherry; PIE-1::GFP embryos used for sorting 1-cell, early and late embryo populations for RNA-seq in Fig. 3a. All graphs show mean of replicates with standard deviation.

**Extended Data Fig. 5:**
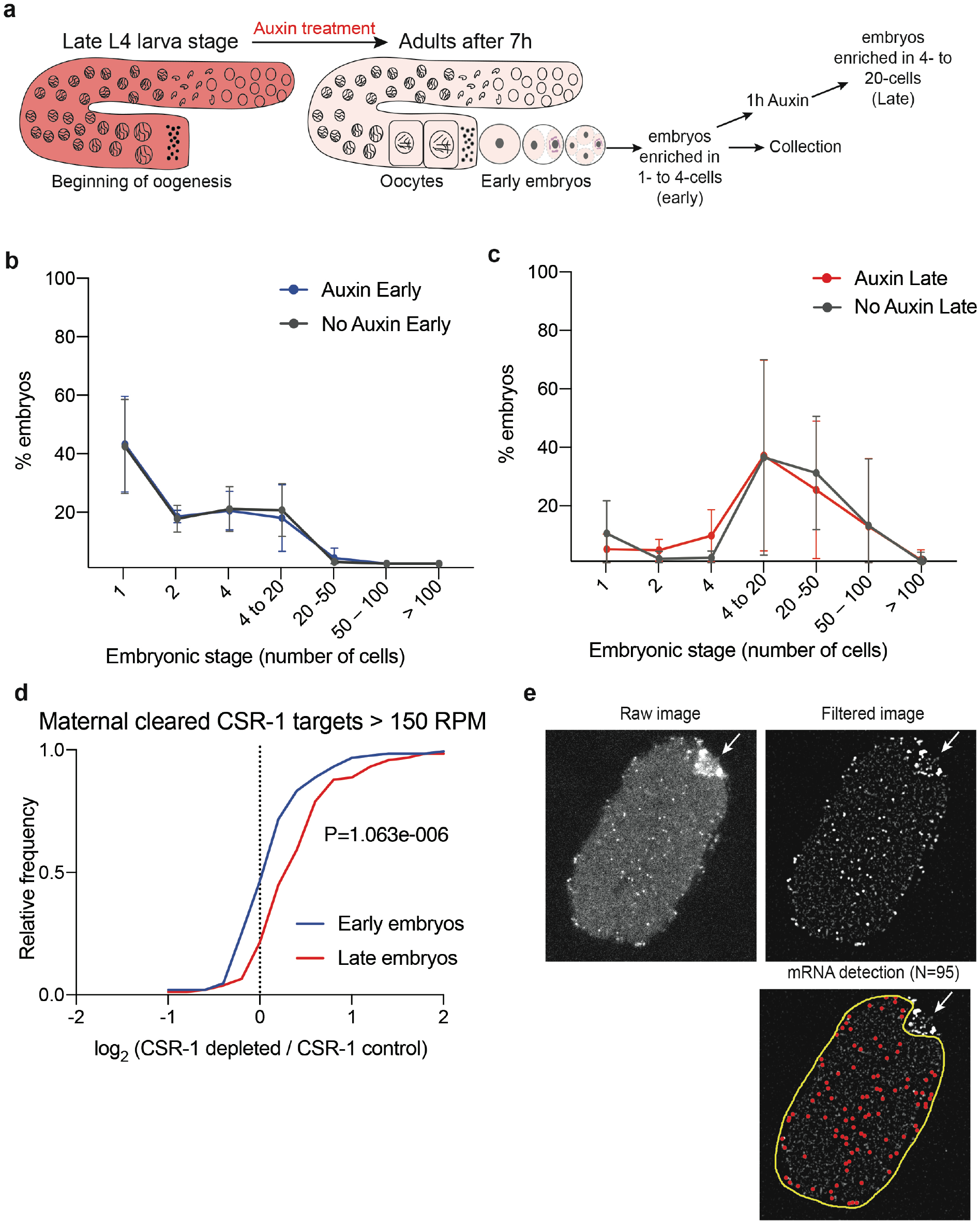
The degradation of maternal mRNAs by CSR-1 occurs in early embryos. **a**, Schematic of strategy to collect embryo preparations at different stages fully depleted of CSR-1. **b**, **c**, Degron::mCherry::3xFLAG::HA::CSR-1 embryos treated with auxin or ethanol (No Auxin) of early **b**, and late embryos **c**, used for the RNA-seq shown in **d**. All graphs show mean of replicates with standard deviation. **d**, Cumulative distribution of the log2 fold change of maternal cleared mRNA targets (> 150 RPM) in CSR-1 depleted early embryos compared to CSR-1 depleted late embryos. Two-tailed P values were calculated using Mann–Whitney–Wilcoxon tests. **e**, Example of quantified image by smFISH of *C01G8.1* mRNA target taken in the central plane of embryos at 20-cells stage. Only somatic cell areas were quantified and germline blastomere were excluded from the quantification. Arrow indicates germline blastomere mRNAs. DAPI staining was used to count number of cells per embryo.

**Extended Data Fig. 6:**
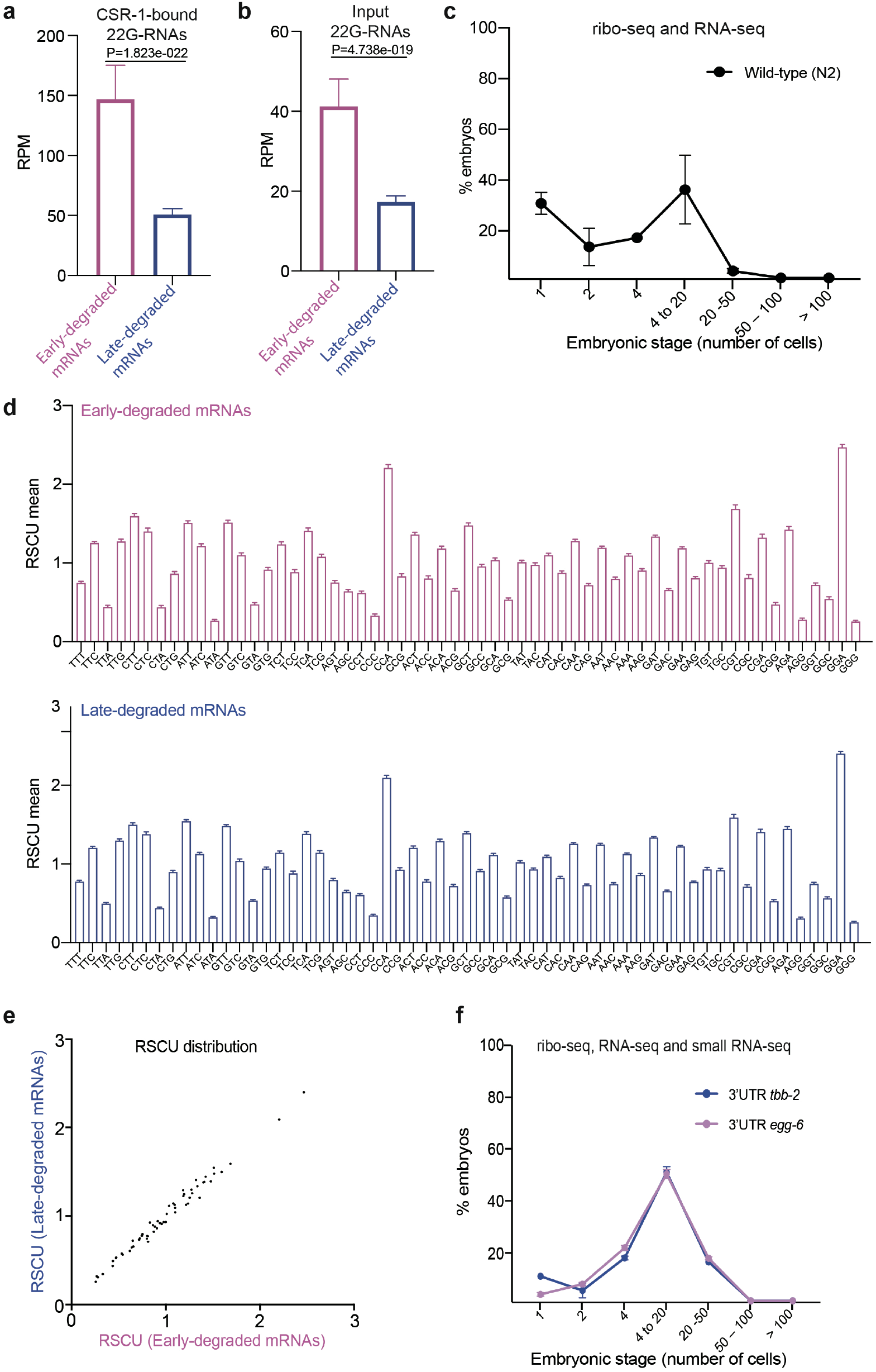
The early and late degraded mRNAs in embryos are differentially regulated by CSR-1. **a**, **b**, Abundance of CSR-1-bound 22G-RNAs (A) or total input 22G-RNAs (B) antisense to early- and late-degraded mRNAs. Median levels and 95 % confidence intervals of normalized 22G-RNA read abundances are shown. Two-tailed p value calculated using the Mann-Whitney-Wilcoxon tests is shown. **c**, *Wild-type* embryos used for Ribo-seq and RNA-seq in **Fig. 4c**, **d**. **d**, Relative synonymous codon usage (RSCU) of early-(top) or late-degraded mRNAs (bottom). Bars show mean with 95% confidential interval. **e**, Average RSCU for each codon in early-or late-degraded mRNAs. **f**, Embryo populations from the two single-copy transgenic reporters expressing *mCherry::h2b* mRNA fused egg-6 3’UTR (early-degraded mRNAs) or tbb-2 3’UTR (late-degraded mRNAs) that have been used for Ribo-seq, RNA-seq and small RNA-seq in **Fig. 5d, e.** All graphs show mean of replicates with standard deviation.

**Extended Data Fig. 7:**
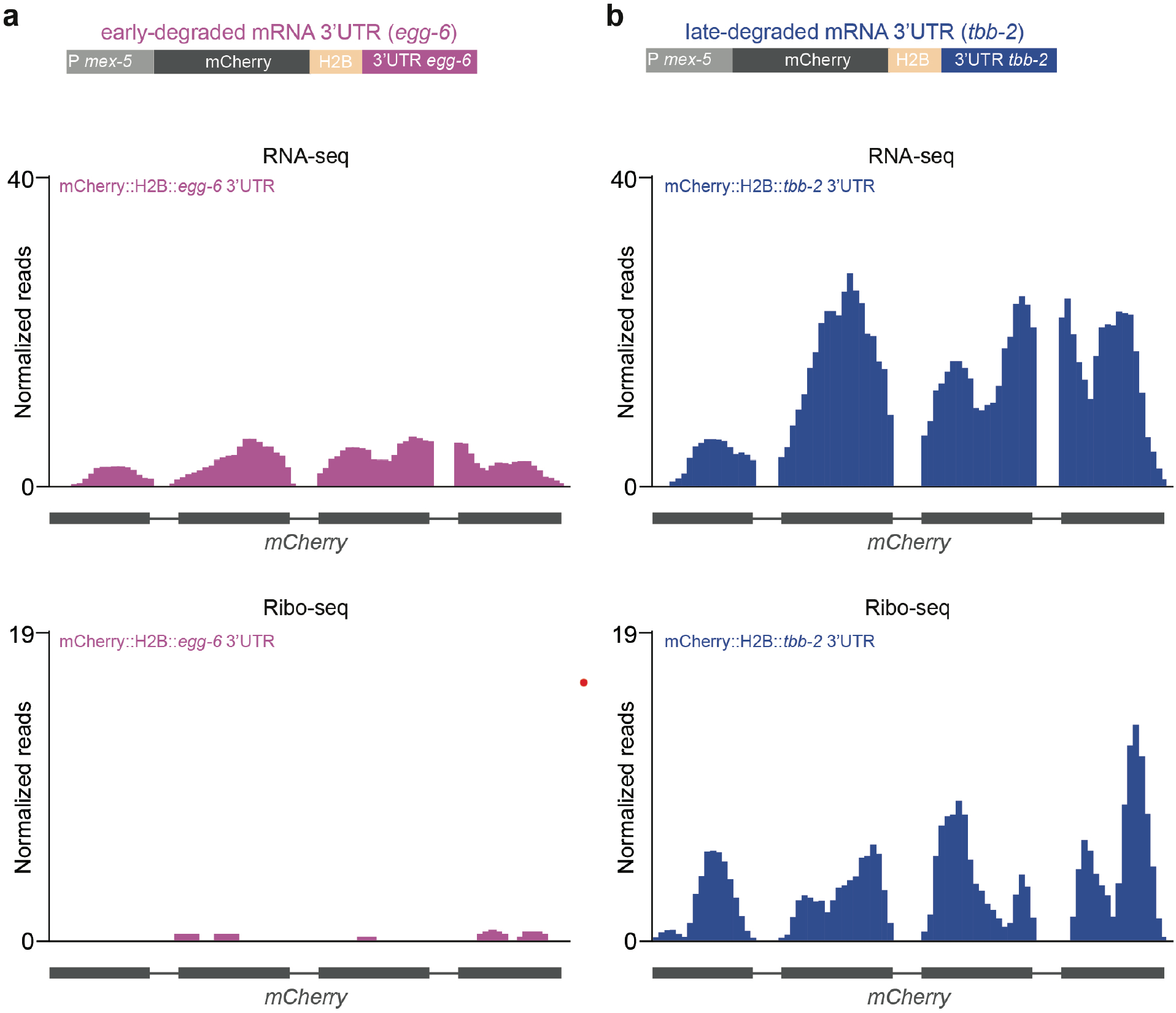
3’UTRs derived from early- or late-degraded mRNAs affects translation and mRNA decay in embryos. Genomic view of normalized reads from RNA-seq or Ribo-seq on *mCherry* coding sequence in embryos expressing *mCherry::h2b* transgene fused to 3’UTR derived from (**a)** early- (*egg-6 3’UTR*) or (**b)** late-degraded (*tbb-2 3’UTR*) mRNAs. The Schematic of the germline-expressed single-copy *mCherry::h2b* transgenic mRNA fused to 3’UTR derived from early- (*egg-6 3’UTR*) (a) or late-degraded (*tbb-2 3’UTR*) (b) mRNAs is shown above.

## SUPPLEMENTAL TABLES

**Supplemental Table 1:** Gene lists generated and used in this study.

**Supplemental Table 2:** Strain lists generated and/or used in this study.

**Supplemental Table 3:** CRISPR-Cas9 guide RNA lists used in this study.

**Supplemental Table 4:** Primer lists used for RT-qPCR.

**Supplemental Table 5**: Oligos used for smFISH.

## METHODS

### *C. elegans* strains and maintenance

Strains were grown at 20 °C using standard methods ^38^. The wild-type reference strain used was Bristol N2. A complete list of strains used in this study is provided in Table S2.

### Generation of transgenic animals

#### Generation of CRISPR–Cas9 lines

CRISPR-Cas9 alleles were generated as described previously ^23^. We used a *mCherry::3×Flag::ha::csr-1* ^23^ as an entry strain to introduce an auxin-inducible Degron tag and obtain a *Degron::mCherry::3xFlag::ha::csr-1* strain suitable for the auxin inducible degradation of CSR-1, after crossing with CA1352, a strain carrying a single copy of the germline expressed TIR-1 protein ^22^. CRISPR-Cas9 guide RNA sequences are listed in Table S3.

#### Generation of MosSCI lines

Strains carrying a single copy insertion of *mex-5P::gfp::csr-1::tbb-2 3’UTR* or *mex-5P::gfp::csr-1(ADH)::tbb-2 3’UTR* were generated by MosSCI ^39^ and the Gateway Compatible Plasmid Toolkit ^39^. The sequences of *csr-1* and *csr-1 (ADH)* were amplified from genomic DNA by PCR from strains carrying either a wild type copy of *csr-1* or *csr-1(ADH)* and cloned in pENTR-D-TOPO (Invitrogen). Plasmids used for Multisite Gateway reaction are listed as follows: pJA245, pCM1.36, pCFJ212. Strain EG6703 was used as an entry strain for single insertion in chromosome IV.

#### 3’UTR replacement experiment

A sequence of 800bp downstream of the stop codon of egg-6 was amplified from genomic DNA by PCR and used as 3’ UTR. We used the single-copy *transgene mCherry::his-11::tbb-2 3’UTR* as entry strain to insert *egg-6 3’UTR* by CRISPR-Cas9 as described above. *egg-6 3’UTR* was inserted right after the STOP codon of *his-11*. The specific expression of the inserted UTR was validated by RT-qPCR.

### Immunostaining

Gonads were dissected on PBS, 0.1 % Tween-20 (PBST) containing 0.3 mM levamisole on 0.01 % poly-lysine slides. Samples were immediately freeze cracked on dry ice for 10 min and fixed at −20°C for 15 min in methanol, and 10 min in acetone. Blocking was performed for 30 minutes at room temperature. Primary antibody was incubated overnight at 4°C in PBS, 0.1 % Tween-20, 5 % BSA. The secondary antibody was incubated for 1 hour at room temperature. Washes were performed with PBS, 0.1 % Tween-20 (PBST). DNA was stained with DAPI. For embryo immunostaining, a drop of embryos from bleached adults was used instead of gonads. The primary antibodies used were anti-Flag (Sigma, F1804) and anti-PRG-1 (gift from the Mello Lab) antibodies at a dilution of 1:500 and 1:800 respectively, and the secondary antibodies used were goat anti-mouse (Invitrogen, Alexa Fluor 488) and goat anti-rabbit (Invitrogen, Alexa Fluor 568) antibodies at a dilution of 1:500.

### Auxin-inducible depletion of CSR-1

Auxin-inducible depletion has been performed as described in ^22^. 250 mM Auxin stock solution was prepared in Ethanol and stored at 4°C. Auxin plates or Ethanol plates were prepared by the addition of Auxin or only Ethanol to NGM plates (final concentration: 500 μM auxin, 0.5 % Ethanol for Auxin plates and 0.5 % Ethanol for Ethanol plates). Plates were seeded with OP50 *E. Coli*, stored at 4°C and warmed at room temperature before the experiment. Worms were placed on Auxin or Ethanol plates from L1 or at the beginning of oogenesis as explained for each experiment.

### Brood size assays

#### WT and CSR-1 mutants

Single L1 larvae were manually picked and placed onto NGM plates seeded with OP50 *E. coli* and grown at 20°C until adulthood and then transferred on a new plate every 24 hours for a total of 2 transfers. The brood size of each worm was calculated by counting the number of embryos and larvae laid on the 3 plates. For the auxin-depleted experiments NGM plates were supplemented with Ethanol (control) or Auxin.

#### Brood size of CSR-1 auxin-depleted worms during oogenesis

For CSR-1 depletion during oogenesis: brood size assay has been performed as described above with the exception that worms were grown from L1 on regular NGM plates and transferred onto Auxin or Ethanol plates 44h after hatching and transferred on new corresponding Auxin and Ethanol plates every 24 hours for a total of 2 transfers.

#### Embryonic lethality assay of CSR-1 rescue experiments

Single L1 larvae of strains carrying Degron::CSR-1 complemented with the single-copy insertion Mex-5P::GFP::CSR-1::tbb-2 3’UTR or Mex-5P::GFP::CSR-1(ADH)::tbb-2 3’UTR were manually picked and placed onto NGM plates seeded with OP50 *E. coli* and grown at 20°C until adulthood. Since transgene was prone to silencing, adult worms were allowed to lay at least 65 embryos and then adults were removed from the plates and imaged to check for GFP::CSR-1 expression. Only plates from GFP expressing adults were used for the assay. The percentage of embryonic lethality is calculated by dividing the number of dead embryos for the total number of laid embryos.

### Collection of early embryos populations

Synchronous populations of worms were grown on NGM plates until adulthood and were carefully monitored using a stereomicroscope and bleached shortly after worms started to produce the first embryos. After bleaching, Early embryos were washed with cold M9 buffer to slow down embryonic development and immediately frozen in dry ice. A small aliquot of embryo pellet (2 μL) was taken right before freezing and mixed with 10 μL VECTASHIELD^®^ Antifade Mounting Medium with DAPI (Vector laboratories) and immediately frozen in dry ice. DAPI stained embryos were defrosted on ice and used for counting cell nuclei and score the embryonic cell stage of the population.

#### Collection of CSR-1 depleted early embryos populations

Early embryos were collected as described above with the exception that worms were transferred on Auxin or Ethanol plates at the beginning of oogenesis.

#### Collection of early and late CSR-1 depleted embryo populations

Early populations of CSR-1 depleted embryos were collected as described above with the exception that harvesting was performed at the very beginning of embryo production to further enrich the population in early staged embryos. After bleaching, an aliquot of the same population was allowed to develop further for 1 to 2h in M9 containing 500 μM Auxin, 0.5 % Ethanol or 0.5 % Ethanol. Embryonic cell stage was scored, and harvesting was performed when embryo populations reached the desired developmental stage.

### Small RNA-seq library preparation

Total RNA from staged embryo preparations with RIN>9 was used to generate small RNA libraries. The library preparation was performed essentially as described previously ^23^.

### RNA IP

A synchronous population of 40,000 worms (48h after hatching) or a preparation of at least 150,000 early embryos was collected and suspended in extraction buffer (50 mM HEPES pH 7.5, 300 mM NaCl, 5 mM MgCl_2_, 10 % glycerol, 0.25 % NP-40, protease inhibitor cocktails (Thermo Scientific), 40 U/mL RiboLock RNase inhibitors (Thermo Scientific)). Samples were crushed in a metal dounce on ice performing at least 40 strokes. Crude protein extracts were centrifuged at 12,000 rpm at 4°C for 10 minutes. Protein was quantified using Pierce™ 660 nm Protein Assay Reagent (Thermo Scientific) and 1 mg (for adults) or 700 μg (for embryos) of protein extract was used for RNA immunoprecipitation as described in ^23^ and used for sRNA-seq library preparation.

### GRO-seq on CSR-1 depleted embryos

Populations containing at least 40,000 CSR-1 depleted early embryos were collected as described above. Early embryos were resuspended in 1.5 mL Nuclei extraction buffer (3 mM CaCl_2_, 2 mM MgCl_2_, 10 mM Tris HCl pH 7.5, 0.25 % Np-40, 10 % Glycerol, Protease inhibitors and RNase inhibitor 4U/mL) and transferred to a steel dounce and stroked 40 times. The lysate was cleared from cell debris by centrifuging at 100×g and nuclei were pellet at 1000×g and washed 4 times with Nuclei extraction buffer. Nuclei were washed once with Freezing buffer (50 mM Tris HCl pH 8, 5 mM MgCl_2_, 0.1 mM EDTA) and resuspended in 100 μL Freezing buffer.

#### Nuclear Run-On reaction and RNA extraction

Nuclear Run-On (NRO) reaction was performed by addition of 100 μL NRO 2x buffer (10 mM Tris HCl, 5 mM MgCl_2_, 1 mM DTT, 300 mM KCl, 1 % Sarkosyl, 0.5 mM ATP, CTP and GTP and 0.8 U/μL RNase inhibitor) and using 1 mM Bio-11-UTP final concentration and incubation for 5 minutes at 30°C. NRO reaction was stopped by the addition of TRIzol LS reagent (Ambion) and RNA extraction was performed following the manufacturer’s instructions. Purified RNA was fragmented by the addition of reverse transcriptase buffer and incubation for 7 minutes at 95°C.

#### Biotin RNA enrichment

Biotinylated nascent RNAs were bound to 30 μL Dynabeads MyOne Streptavidin C1 (Invitrogen and) washed 3 times as described in ^40^ and purified with TRIZOL reagent.

#### RNA 5’-end repair

5’-OH or fragmented RNAs were repaired using Polynucleotide kinase (Thermo scientific) following manufacturer instructions and incubated at 37°C for 30 minutes. RNA was purified with Phenol:Chloroform and precipitated by the addition of 3 volumes of Ethanol, 1/10^th^ volumes of 3 M Sodium Acetate and 30 μg Glycoblue coprecipitant (Ambion).

#### Ligation of 3’ and 5’ Adapter Oligos and 2^nd^ and 3^rd^ Biotin enrichments

RNA was ligated to 3’ end adapter using T4 RNA ligase 2 Truncated KQ (home-made) for 16h at 15°C. After ligation RNA was purified using solid-phase reversible immobilization beads (SPRI beads) and biotinylated RNA was enriched as described above. After purification RNA was ligated at 5’ end using T4 RNA ligase 1 for 2 h at 25°C. RNA was purified using SPRI beads and biotinylated RNA was enriched for a third time as described above.

#### Reverse transcription and amplification of cDNA libraries

Purified RNA was reverse transcribed using SuperScript IV Reverse Transcriptase (Thermo Fisher Scientific) following manufacturer conditions except that reaction was incubated for 1h at 50°C. cDNA was PCR amplified with specific primers using Phusion High fidelity PCR master mix 2x (New England Biolab) for 18-20 cycles and sequenced on Illumina Next 500 system.

### Sorting of *C. elegans* embryos

Sorting of *C. elegans* embryos was performed as described in ^28^ with the following modifications: a strain expressing both mCherry::OMA-1 and PIE-1::GFP was used to collect embryos at three different developmental stages (see **Extended Data Fig. 3c**); embryos after bleaching were fixed with 2% formaldehyde in M9 to block cell division. After sorting, embryos were reverse crosslinked in 250 μL RIPA buffer with RNase inhibitors for 30 minutes at 70°C and RNA was extracted with TRIzol LS (Ambion) following manufacturer instructions.

### Strand-specific RNA-seq library preparation

DNase-treated total RNA with RIN > 8 was used to prepare strand-specific RNA libraries. Strand-specific RNA-seq libraries were prepared as described previously ^23^.

### RT-qPCR

1 μg DNase treated total RNA was used as a template for cDNA synthesis using random hexamers and M-MLV reverse transcriptase. qPCR reaction was performed using Applied Biosystems Power up SYBR Green PCR Master mix following the manufacturer’s instructions and using an Applied Biosystems QuantStudio 3 Real-Time PCR System. Primers used for qPCR are listed in Table S4.

### Single-molecule FISH (smFISH)

Single-molecule FISH (smFISH) was performed as described in ^41^. Briefly, embryos were harvested by bleaching, immediately resuspended in Methanol at −20°C, freeze cracked in liquid nitrogen for 1 minute and incubated at −20°C overnight. Embryos were washed once in wash buffer (10 % formamide, 2x SSC buffer (Ambion)) and hybridized with the corresponding FLAP containing probes in 100 μL hybridization buffer (10 % dextran sulfate, 2 mM vanadyl-ribonucleoside complex, 0.02 % RNAse-free BSA, 50 μg *E. coli* tRNA, 2× SSC, 10 % formamide) at 30°C overnight. Hybridized embryos were washed twice with wash buffer and once in 2x SSC buffer before imaging. Right before imaging, embryos were resuspended in 100 μL anti-fade buffer (0.4 % glucose, 10 μM Tris-HCl pH8, 2x SSC) with 1 μL catalase (Sigma-Aldrich) and 1 μL glucose oxidase (3.7 mg/mL, Sigma-Aldrich) and stained with VECTASHIELD^®^ Antifade Mounting Medium with DAPI (Vector laboratories). DAPI staining was used to select embryos at 20-cell stage. Images of the central plane of embryos at desired developmental stages were acquired on a Zeiss LSM 700 confocal microscopy. Oligos used for smFISH of *C01G8.1* mRNA target are listed in Table S5.

The detection of single mRNA molecules was performed with the open-source Matlab package FISH-quant as previously described ^42^. Briefly, RNA signal was enhanced by a 2-step convolution of the image with Gaussian Kernels. First, the image background obtained by convolution with a large Gaussian Kernel was estimated and then subtracted. Second, the resulting image was filtered with a small Gaussian Kernel to further enhance the signal-to-noise ratio. RNA spots were detected with a local maximum algorithm. For each embryo, a manually drawn outline was used to limit the detection to the somatic blastomeres and exclude the germline blastomere.

### Ribo-seq

Ribo-seq has been performed as described in ^43^ with some modifications. Briefly, embryos harvested by bleaching were lysed by freeze grinding in liquid nitrogen in Polysome buffer (20 mM Tris-HCl pH 8, 140 mM KCl, 5 mM MgCl_2_, 1 % Triton X-100, 0.1 mg/mL cycloheximide) and 1 mg extract was digested by RNase I (100 U) at 37°C for 5 min and then fractionated on sucrose gradient (10-50 %) by ultracentrifugation at 39,000 rpm in a SW41-Ti rotor (Beckman coulter). RNA from monosome fraction was DNase treated and fragments of 28-30 nucleotides were size selected after running on a 15 % TBE-Urea gel. 28-30 nucleotide Ribosome protected fragments (RPF) were cloned with the previously described sRNA-seq library preparation approach with the following modifications: 3’phosphate was removed and 5’ end was phosphorylated by treating RNA with Polynucleotide kinase.

### Sequencing data analyses

For Ribo-seq data, The 3′ adapter was trimmed from raw reads using Cutadapt ^44^ v.1.18 using the following parameter: -a TGGAATTCTCGGGTGCCAAGG –discard-untrimmed. Trimmed reads of size ranging from 28 to 30 nucleotides were selected using bioawk (https://github.com/lh3/bioawk). The selected 28–30-nucleotide reads were aligned to the *C. elegans* genome sequence (ce11, *C. elegans* Sequencing Consortium WBcel235) using Bowtie2 ^45^ v.2.3.4.3 with the following parameters: -L 6 -i S,1,0.8 -N 0. Reads mapping on sense orientation on annotated protein coding genes were considered as Ribosome-Protected Fragments (RPF). Such reads were extracted from mapping results using samtools 1.9 ^46^ and bedtools v2.27.1 ^47^ and re-mapped on the genome.

Analysis for RNA-seq, sRNA-seq, and GRO-seq have been performed as previously described ^23^. Metaprofiles were generated using RPM from sRNA-seq analysis by summarizing normalized coverage information (taken from bigwig files and averaged across replicates) along early degraded targets or late degraded targets using deeptools ^48^. For RNA-seq, GRO-seq and Ribo-seq, read counts associated to annotated genes were obtained from the alignments using featureCounts ^49^ v.1.6.3 and annotations obtained as previously described (*17*). Normalized abundances were estimated from read counts by first normalizing by transcript union exon length (reads per kilobase, RPK), then by million total RPK (transcripts per million, TPM). Mean TPM was computed by taking the mean across replicates. For Ribo-seq data, translational efficiency (TE) was measured as the log_2_ of the ratio between mean TPM of ribosomal protected fragments and of transcripts from RNA-seq, adding a pseudo-count of 1 to both terms of the ratio (in order to avoid discarding genes for which no transcript was detected in RNA-seq).

### RSCU calculation

RSCU for earlier and later degraded mRNAs was calculated using CAI calculator (http://genomes.urv.es/CAIcal/).

### Gene lists

Gene lists are provided in Table S1.

